# Antibiotic production reduces the cost of resource cheaters in *Streptomyces coelicolor*

**DOI:** 10.1101/2025.11.28.691120

**Authors:** Linus Theinert, David M. Norte, Britt Veugelers, Linus Veit, Luis Alfredo Avitia-Dominguez, Daniel E. Rozen

**Author notes:** Joint First Authors (Linus Theinert, David M. Norte). Linus Theinert -, David Norte -, Britt Veugelers - Linus Veit -, Luis Alfredo Avitia-Dominguez - Daniel Rozen.

## Abstract

Soil is a competitive environment containing a variety of resources used for bacterial growth. Complex polysaccharides, like chitin or starch, require the secretion of enzymes that degrade these resources into smaller units before they can be consumed. However, exoenzymes and the products they create are public goods, meaning they can be used by competitors, called “cheaters”, who benefit from public goods even if they do not produce enzymes themselves. Here, we test the hypothesis that antibiotics produced by *Streptomyces* are used to privatize public goods by restricting access to resource cheaters. Using experiments with *Streptomyces coelicolor* and *Bacillus subtilis*, we first show that *B. subtilis* cheating significantly reduces *S. coelicolor* fitness on complex medium (starch) but not on a simple carbon source (maltose) which does not require exoenzyme secretion. Next, we show that antibiotics produced by *S. coelicolor* markedly increase fitness against resource cheaters, despite evidence that antibiotic production is metabolically costly. Finally, we find that the benefits of antibiotic production and the costs of resource cheating are both higher during growth on lower resource concentrations. Our results provide novel insights into the context-dependent costs and benefits of antibiotic secretion in *Streptomyces* and highlight the role of resource complexity and concentration in mediating competitive strategies in bacteria.

## Introduction

Soil is a complex environment where microbes and other microfauna compete for access to resources and space. One of the central axes of microbial competition in soil is for limiting nutrients, which can broadly be partitioned into simple or complex resources (Wang & Kuzyakov, 2024). Whereas simple carbon sources, including sugars like glucose, can be directly transported and used by most bacterial species, complex carbon sources, like starch and chitin, require enzymes to degrade them extracellularly into simpler monomers or oligomers before they can be used for nutrition. Complex polysaccharides are among the most abundant sources of stored carbon in nature, and exoenzyme secretion by soil saprotrophs is crucial for their mineralization (Chater et al., 2010; Sichert & Cordero, 2021). However, while enzyme secretion is essential for growth, it also comes with the associated risk of exploitation (Smith & Schuster, 2019). Exoenzymes are secreted extracellularly, and the products of degradation become freely available to other species, whether they secrete exoenzymes or not. Accordingly, exoenzymes are a type of public good that can provide collective benefits to producing species, but are open to exploitation by “cheaters” that benefit from production while avoiding the costs of production (Allison, 2005). Such dynamics are well studied in terrestrial and marine environments and give rise to trophic cascades where primary degraders act as keystone species for the initial breakdown of insoluble resources (Cordero & Datta, 2016; Dundore-Arias et al., 2019; McClure et al., 2022). However, it is less well understood how these keystone species protect their investment. Here, we test the hypothesis that antibiotics made by streptomycetes can be used to privatize public goods and thereby reduce the costs of resource exploitation by heterotrophic competitors.

Streptomycetes are Gram-positive filamentous saprotrophs that encode and secrete a large repertoire of hydrolytic enzymes that degrade complex carbohydrates like starch, chitin, cellulose, and xylan into their respective monomers and oligomers (Chater et al., 2010; Vionis et al., 1998). Although the composition of exoenzymes varies between species, likely dependent on local features of the environment in which they live, all species encode hundreds of secreted enzymes that play key roles in their biology as well as their interactions with competitors. Several studies have shown that *Streptomyces* species are enriched following cellulose, lignin, or chitin amendment to agricultural fields (Dundore-Arias et al., 2019, 2020; D. Schlatter et al., 2009), suggesting that these recalcitrant resources benefit *Streptomyces* growth. Furthermore, recent results have found that *S. venezuelae* can subsist solely on insect cuticles (Meij et al., 2025). In addition, removing *Streptomyces* species from synthetic bacterial communities grown on chitin strongly limits the growth of secondary consumers. These results confirm that the byproducts of primary degradation can be consumed (exploited) by other competitors (McClure et al., 2022), raising questions about the mechanisms these bacteria use to privatize these resources.

In addition to exoenzymes, streptomycetes are prolific producers of antimicrobial compounds, accounting for the majority of all known antibiotics (Morin et al., 2025). Many functions have been proposed for antibiotics, ranging from intercellular signals to weapons (Jauri et al., 2013; Romero et al., 2011). However, most evidence is consistent with their use in interference competition to kill or inhibit competitors, to provide defense against predators or bacteriophages (Morin et al., 2025; D. C. Schlatter & Kinkel, 2014). Related to this are results showing that antibiotic production is triggered by cues produced by competing species, consistent with the idea that antibiotics are facultatively deployed via “competition sensing” (Abrudan et al., 2015; Traxler et al., 2013; Westhoff et al., 2021). Importantly, however, most experiments that evaluate conditions for antibiotic induction do not test the role or fitness effects of these compounds in direct pairwise interactions. Thus, while it is known that purified antibiotics can inhibit other bacteria in isolation, much as they do during therapeutic use, it is often unclear how these same compounds mediate competitive dynamics (Cornforth & Foster, 2013). Moreover, even in cases where competitive interactions are examined, they are most often conducted in environments with abundant labile resources and between species with equivalent growth rates and resource use (Westhoff et al., 2020). It is therefore relatively unexplored whether the benefits of antibiotics depend on carbon source complexity or the presence of competitors that can exploit exoenzyme production.

In this paper, we use the model species *Streptomyces coelicolor* to study the benefits of antibiotic production when strains are challenged with public goods cheaters. *S. coelicolor* produces several antibiotics whose synthesis coincides with the developmental transition to sporulation (Heul et al., 2018). The temporal association between antibiotic production and development has led to the idea that the two behaviors are functionally coupled, first by driving developmentally regulated programmed cell death (Tenconi et al., 2018) and second to inhibit competing species that can grow on the resources released by lysing cells (Filippova & Vinogradova, 2017). Here, we propose an alternative hypothesis: the benefits of antibiotics in this species are contingent on resource type and concentration and the presence of exploitative competitors. To test this, we studied ecological interactions between *S. coelicolor* and the common soil bacterium *B. subtilis*. To distinguish the role of antibiotics and exoenzyme exploitation, respectively, we used strains of *S. coelicolor* that vary in the production of actinorhodin, an antibiotic that kills *B. subtilis*, and strains of *B. subtilis* that differ in their ability to produce amylase needed to degrade starch. In brief, our results confirm that resource exploitation is extremely costly to *S. coelicolor* and that the ability to mitigate these costs with antibiotics differs in complex and simple resource environments. We also show that the benefits of antibiotics scale with resource concentration. Our results show the importance of both resource type and amounts in mediating the outcome of microbial competition and highlight the context-dependent benefits of antibiotic production in *S. coelicolor*.

## Methods and Materials

### Bacterial strains and culturing conditions

To study the effects of antibiotic production in *S. coelicolor*, we used two strains that differ in their ability to produce actinorhodin: a wild-type strain (*S. coelicolor* M145 A3(2), designated StrepWt) and an otherwise isogenic strain with a mutation in a key activator of actinorhodin synthesis (*S. coelicolor* A3(2) M145 Δ*act*II-ORF4, designated StrepΔact) (Floriano & Bibb, 1996). Both strains were modified to express GFP constitutively and can therefore be quantified using cell sorting, as described below. We used *S. coelicolor* A3(2) M145 Δ*act*II-ORF4Δ*redD* (designated StrepΔactΔred), which is unable to produce actinorhodin and undecylprodigiosin, to study the effects of amylase exploitation during growth on starch medium (Floriano & Bibb, 1996). *S. coelicolor* was routinely grown at 30 °C on Soy Flour Mannitol Agar (SFM) containing per liter: 20 g Soy Flour (DO-IT BV, Barneveld, The Netherlands), 20 g Mannitol (Duchefa Biochemie, Haarlem, The Netherlands), and 15 g agar (Hispanagar, Burgos, Spain) adjusted to pH 7.2. High-density spore stocks were generated by plating 100 μL of spore solution on SFM (Westhoff et al., 2020). After 3-5 days of growth, a cotton disk soaked in 3 mL of 30% glycerol was added to the plates. Spores were extracted by gently rubbing the cotton disk over the confluent lawn of spores and filtering the liquid through the cotton filter to remove the vegetative mycelium. Spore stocks were titered via serial dilution and stored at -20 °C.

To study the impact of resource exploitation, we used two *Bacillus subtilis* strains that vary in the ability to produce amylase, the enzyme needed to degrade extracellular starch: *B. subtilis* P5_B1 (designated BacWt) and *B. subtilis* P5_B1 Δ*amyE* (designated Bac*ΔamyE*), both of which constitutively express mKate (Thérien et al., 2020; van Gestel et al., 2014). Strains were kindly provided by Ákos Kovács (University of Leiden, NL). Frozen culture stocks were prepared by growing strains overnight in 25 mL LB broth at 37 °C under constant shaking (200 rpm), after which they were stored at -20 °C in 20% glycerol.

Modified ISP4 was used for all competition and growth experiments, consisting, per liter, of 1 g MgSO4 (Merck KGaA, Darmstadt, Germany), 1 g NaCl (Carl Roth GmbH + Co. KG, Karlsruhe, Germany), 2 g (NH4)2SO4, 2 g CaCO3 (BDH Chemicals Ltd, Poole, England), 1g K2HPO4 (Avantor Performance Materials Poland S.A., Gilwice, Poland), 20 g of agar (Sphaero Q, Gorinchem, The Netherlands). 1 mL from a stock trace salts solution (0.1 g FeSO4, 0.1 g MnCl2, 0.1 g ZnSO4, 100 mL dH_2_O) was added after autoclaving and pH was adjusted to 7.2. Maltose and soluble starch were prepared as 30% stock solutions in demi water and added to the autoclaved ISP4 base to obtain either 0.1% or 1% maltose or starch ISP4 medium.

### Growth during co-cultures of Streptomyces and Bacillus

To determine if *B. subtilis* growth on starch medium was influenced by proximity to *S. coelicolor* (via starch degradation), we inoculated droplets of ∼10^4^ spores of StrepΔactΔred onto ISP4 plates with 1% starch and incubated the plates overnight at 30°C. The resulting colonies were covered with 4 mL of 0.7% soft water agar containing 100 µL (at ∼10^7^/mL) of each *Bacillus* strain (two replicates each). Images were taken with a stereo microscope after 24 hours of further incubation. 100 randomly selected colonies from each plate were measured for size and distance from the *Streptomyces* colony using the oval tool in Fiji.

### Fitness effects of antibiotic production and resource exploitation

Competition assays between bacterial strains were carried out on ISP4 plates containing 0.1%, 1% starch, or 1% maltose. We examined four combinations: StrepWt vs BacilWt, StrepWt vs Bacil*ΔamyE*, StrepΔact vs BacilWt, and StrepΔact vs Bacil*ΔamyE*. Strains were differentiated using their expression of either GFP (*S. coelicolor*) or mKate (*B. subtilis*). This allowed us to quantify the frequencies of each competitor using a BioRad S3e Cell Sorter flow cytometer (Bio-Rad Laboratories, USA). The sorter was configured for dual-channel fluorescence detection (GFP: 488 nm excitation, 525/30 nm emission; mKate: 561 nm excitation, 615/25 nm emission). Monoculture controls of each fluorophore were used to define gating boundaries and correct for spectral overlap. Events were gated on forward and side scatter to select for spores and single cells and exclude debris or aggregates.

Experiments were initiated by mixing ∼ 10^6^/mL CFU equal parts of either *S. coelicolor* strain and a corresponding *B. subtilis* strain. This was analyzed using the cell sorter to obtain initial frequency values. Replicate plates were inoculated by uniformly spreading 100 µL of the mix using glass beads onto each medium type, after which they were incubated for 7 days at 30°C. Spores were collected by covering the surface of the plate with 3 ml of 30% glycerol, gently rubbing a cotton disk over the lawn to dislodge the spores, and filtering the liquid through a cotton filter to remove the vegetative mycelium. The filtrate was diluted 100x, and an aliquot was run through the sorter to obtain final frequency values. Fitness (f) was quantified following (Jiricny et al., 2010)

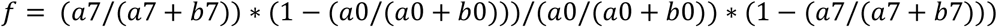

where *a*0 and *a*7 are the *S. coelicolor* strain at start and finish, respectively, and *b*0 and *b*7 are the *B. subtilis* strains at start and finish, respectively.

### Streptomyces growth rates

*Streptomyces* growth rate as a function of carbon source concentration was quantified by measuring radial growth via time-lapse imaging of replicate colonies. 10 µL containing 50 spores of StrepWt or StrepΔact were spread on modified ISP4 containing either 0.1% or 1% starch. Plates were incubated at 30°C in a Reshape Imaging Device (Reshape Biotech, Copenhagen, Denmark) for 7 days. For each strain and starch concentration, two replicate plates were imaged every hour, allowing over 60 isolated colonies per plate to have their area calculated every hour using the Discovery platform of Reshape Biotech. Each colony was measured from the first instance of detection until 7 days after incubation started. Finally, hourly measurements of colony area were converted into average hourly area growth speed (*mm*^2^/*h*) of each colony.

### Streptomyces spore production on starch medium

*Streptomyces* spore production as a function of carbon source concentration was quantified by measuring total CFU from single colonies. 10 µL containing 50 spores of *Strep*Wt or StrepΔact were spread onto modified ISP4 containing either 0.1% or 1% starch. Plates were incubated at 30°C for 7 days. For each strain and medium, six isolated single colonies were scraped from the agar, dropped into tubes containing 1 mL of 30% glycerol, and thoroughly vortexed for 10 minutes. Spores were separated from mycelia using a custom microfilter consisting of a cotton wool-plugged 200 μL pipette tip inserted into a hole drilled into the lid of a sterile 1.5 mL centrifuge tube (Avitia Domínguez et al. 2025). 200 μL of the colony-suspended solution was added into the microfilter and centrifuged at a speed of 8000 rpm for 2 min. The filtrate was serially diluted and plated onto SFM agar plates to determine the spore output of each assayed colony.

### Quantifying antibiotic production

To quantify antibiotic production under different growth conditions, we measured the halo of inhibition of a target *Bacillus* strain grown atop *S. coelicolor* colonies. *Streptomyces* colonies were established by plating 2 µL drops containing 10^5^ *S. coelicolor* Wt spores onto modified ISP4 containing 1% or 0.1% starch. After five days of incubation, overnight cultures of *B. subtilis* Wt were diluted to roughly 10^8^/mL after which 100 µL was mixed with 4 mL molten LB soft agar (0.7%), and poured over the inoculated plates to create a uniform overlay lawn of susceptible bacteria. After 24 hours of incubation, the plates were imaged with a Zeiss Axiozoom v16 stereo microscope. To calculate the halo of inhibition caused by antibiotic production, the area was manually measured in Fiji using the circle tool.

### Statistical analysis

Statistical comparisons between strains and treatment conditions were analyzed with one- and two-way ANOVAs using the aov function in R version 4.5.1 and R Studio version 2025.09.2. Distance-dependent growth in the presence of *S. coelicolor* on starch medium was analyzed using linear regression using the lm function in the same software.

## Results

### BacΔamyE obligately exploits S. coelicolor on starch medium

To confirm that Bac*ΔamyE* was unable to grow on starch-supplemented medium, we compared the growth of this strain on MM with either 1% starch or 1% maltose. As expected, we observed rapid growth to high densities on maltose and negligible growth on starch (Supplementary Figure 1). Next, we tested if the growth of Bac*ΔamyE* could be restored on starch-supplemented ISP4 medium when grown adjacent to StrepΔactΔred, where it could consume resources produced by starch degradation. We used an antibiotic-deficient *S. coelicolor* strain to allow *B. subtilis* to grow as close to its competitor as possible without risk of inhibition. Preliminary results confirmed that neither StrepΔact nor StrepΔactΔred inhibit *B. subtilis* growth, indicating that undecylprodigiosin production has no discernible influence on antibiotic-driven competitive advantages (Supplementary Figure 2). While the growth of *B. subtilis* Wt under these conditions was independent of the distance to the *S. coelicolor* colony (*R*^2^= 0.02097 p = 0.041), Bac*ΔamyE* colony size declined rapidly with increasing distance (Figure 1, Linear regression *R*^2^= 0.3321 p < 0.001). These results confirmed that while growth of BacWt is independent of amylase secretion by *S. coelicolor*, Bac*ΔamyE* is dependent on resources provided by *S. coelicolor* amylase production for growth.

**Figure 1.**
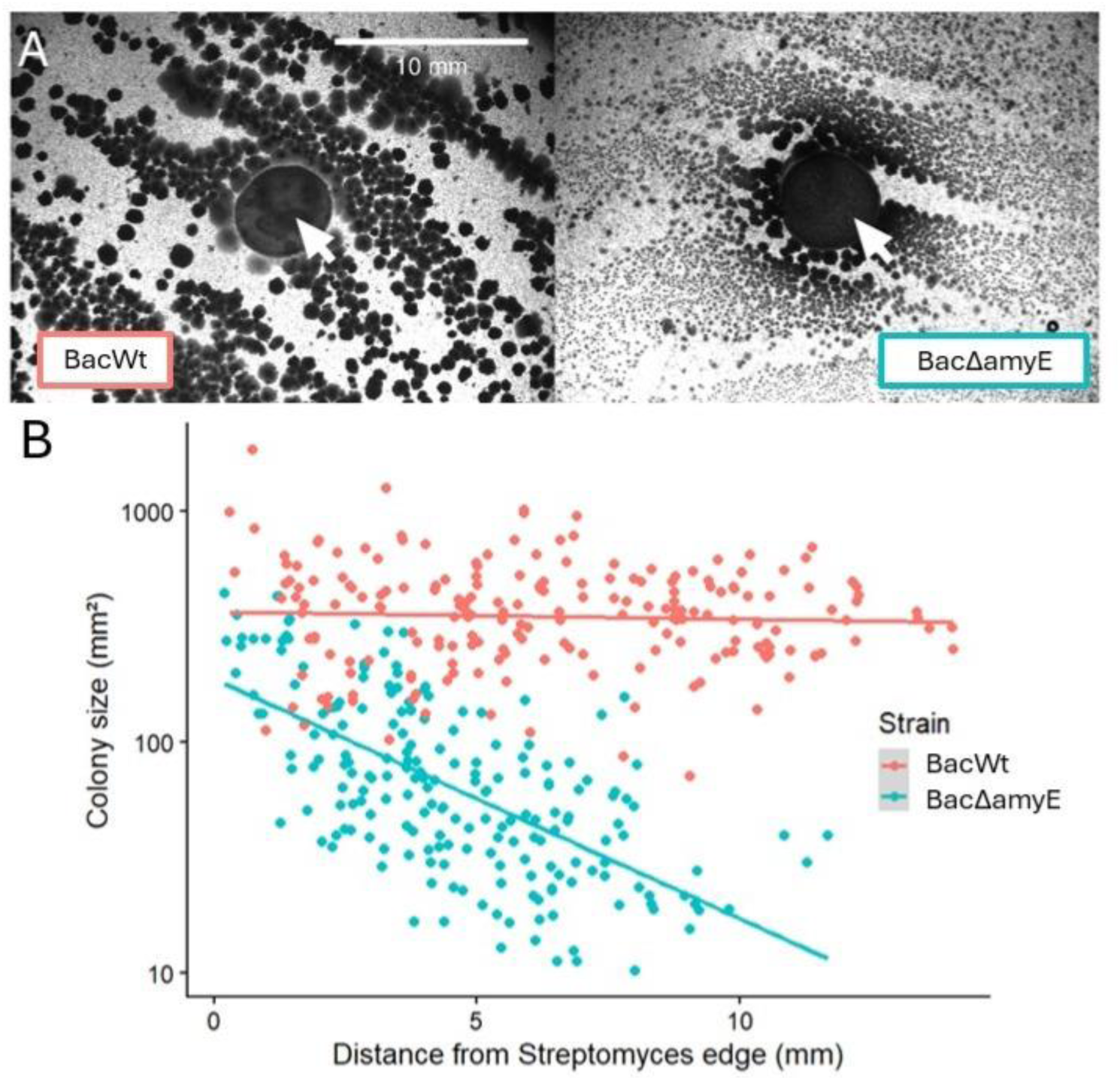
B. subtilis ΔamyE (BacΔamyE) and Wt (BacWt) growth in the presence of S. coelicolor ΔactΔred (StrepΔactΔred) on 1% starch medium. A) BacΔamyE colonies exploit Streptomyces starch degradation for growth. White arrows indicate the StrepΔactΔred macrocolony. B) BacΔamyE growth on starch is dependent on proximity to StrepΔactΔred, unlike BacWt.

### Antibiotic production provides medium-dependent benefits against wild-type and exploiter strains of B. subtilis

After confirming that Bac*ΔamyE* could exploit *S. coelicolor*, we next used pairwise competition assays under different growth conditions to investigate the fitness costs of exploitation and whether these costs were mitigated by antibiotic production. The results of these assays are shown in Figure 2 and lead to several conclusions. First, regardless of the ability to produce antibiotics or the identity of the *B. subtilis* competitor, there are dramatic differences in the fitness of *S. coelicolor* on the complex starch medium compared to its fitness on maltose. While *S. coelicolor* outcompetes *B. subtilis* on starch, it loses comprehensively on maltose (ANOVA 2-way: F_1,10_ = 102.72, p < 0.001). Importantly, this fitness difference is not caused by growth deficits of *S. coelicolor* on maltose, where its growth in isolation is even faster than its growth on starch (Supplemental Figure 3). Second, while there are significant costs of resource exploitation to *S. coelicolor* on starch medium (ANOVA 1-way: F_1,10_ = 18.84, p < 0.001), resulting in a 3-fold decline in fitness when comparing StrepWT vs BacWT to competition against Bac*ΔamyE*, these are not observed during growth on maltose (ANOVA 1-way: F_1,10_ = 0.02, p = 0.997), likely because *B. subtilis* growth is independent of *S. coelicolor* on this medium. Third, we found significant benefits of antibiotic production on both media types (ANOVA 1-way: Starch F_1,10_ = 21.09, p < 0.001; Maltose F_1,10_ = 36.28, p < 0.001). However, these benefits are far larger on starch than on maltose (ANOVA 2-way Interaction, F_1,20_ = 36.02, p < 0.001), where loss of production led to a 9-fold decline in fitness on starch and a 2-fold cost on maltose. Notably, even though antibiotics benefit *S. coelicolor* on maltose, this negligible benefit is unable to overcome the competitive differences between the two species on this carbon source. Finally, we found no differences in the fitness of StrepΔact when competing versus either *B. subtilis* strain on either medium, suggesting that there is a limited added cost of resource exploitation (ANOVA 2-way, F_1,20_ = 0.189, p = 0.669) in the absence of antibiotic production. Altogether, these results show that resource exploitation imparts significant fitness costs that are media-dependent, and that these costs can be mitigated through antibiotic production.

**Figure 2.**
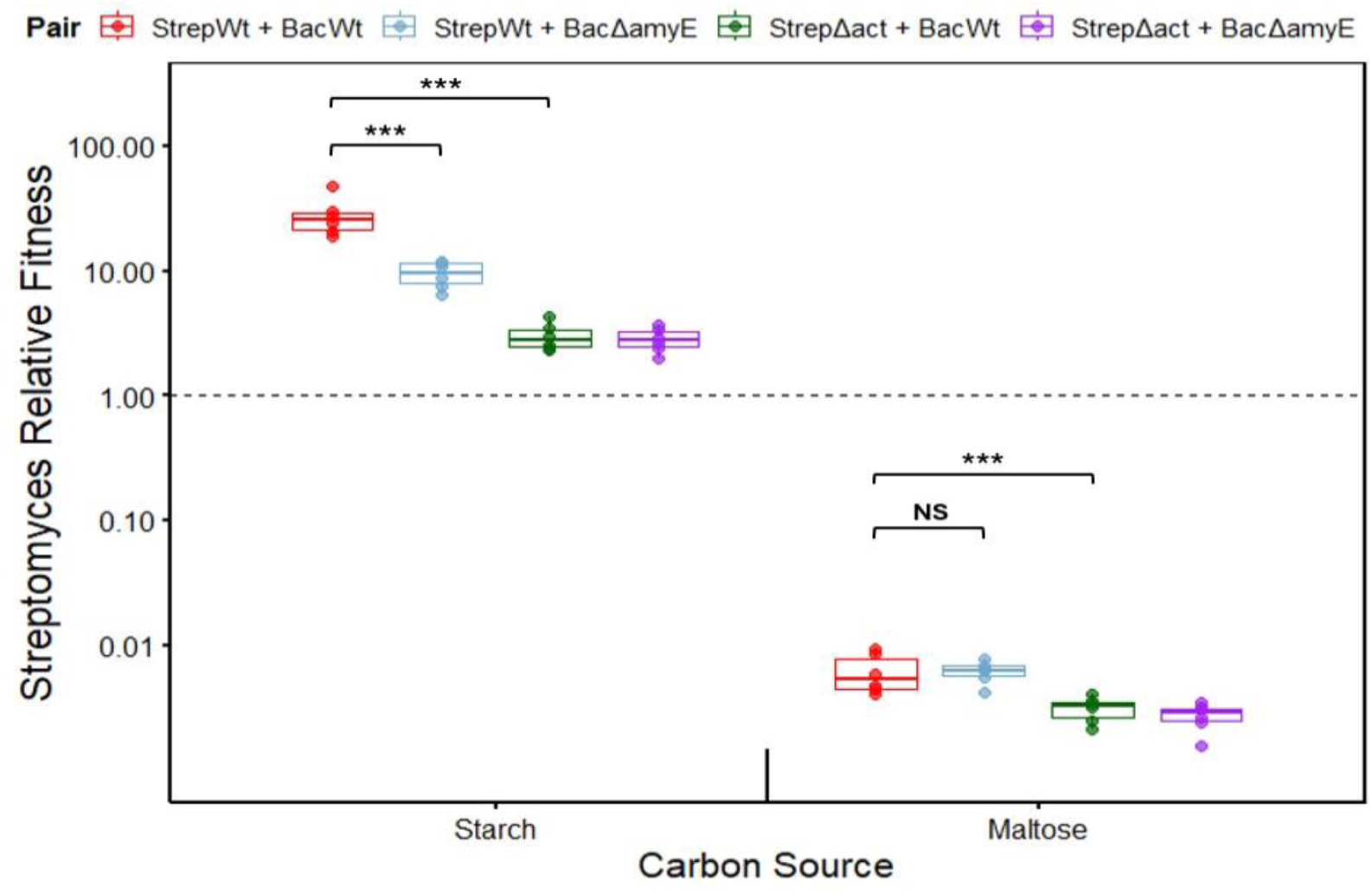
Fitness of *S. coelicolor* in competition with *B. subtilis* on starch and maltose. While BacΔamyE has a detrimental effect on StrepWt fitness on starch, this is absent on maltose. Antibiotic production provides a fitness benefit in both environments and overcomes the cost of cheating by BacΔamyE. *** corresponds to p < 0.001.

### The costs and benefits of antibiotic production and exploitation are resource concentration-dependent

Previous studies have suggested that the costs and benefits of public goods are influenced by resource concentration (Brockhurst et al., 2008). This prompted us to test whether the benefits of antibiotic production and the costs of resource exploitation would vary as a function of starch concentration. As in the previous experiments, we established competition assays between the four possible combinations of *S. coelicolor* and *B. subtilis* strains, and then measured *Streptomyces* fitness as a function of whether strains were competed in either 0.1% or 1% starch (Figure 3). Overall, starch concentration lead to significant differences in the fitness of different pairings (ANOVA 2-way Interaction, F_3,40_ = 47.68, p < 0.001). We found that the fitness of StrepWT was significantly higher on lower starch concentrations (ANOVA 1-way: F_1,22_ = 14.21, p < 0.001). Although there is a significant cost of resource exploitation in both concentrations (ANOVA 1-way: starch 1% F_1,10_ = 26.3, p < 0.001; starch 0.1% F_1,10_= 52.35, p < 0.001), the magnitude of the cost varies between the two concentrations (ANOVA 2-way Interaction: F_1,20_ = 31, p < 0.001), with higher fitness costs on lower (3.7-fold decline on 0.1%) than on higher starch concentrations (2.5-fold decline on 1%). Similar to results in Figure 2, we found no differences in the fitness of StrepΔact when competing versus either *B. subtilis* strain, regardless of concentration (ANOVA 2-way, F_1,20_ = 0.467, p = 0.2268). Finally, while antibiotics are highly beneficial under both conditions (ANOVA 2-way: F_1,20_ = 140.6, p < 0.001), the magnitude of the benefit is markedly higher in low starch (ANOVA 2-way Interaction: F_1,20_ = 56.85, p < 0.001). While the loss of antibiotic production leads to a 52-fold fitness reduction in 0.1% starch, this is only 9-fold in 1% starch, suggesting that *S. coelicolor* is more effective at protecting against resource exploitation on lower resource concentrations.

**Figure 3.**
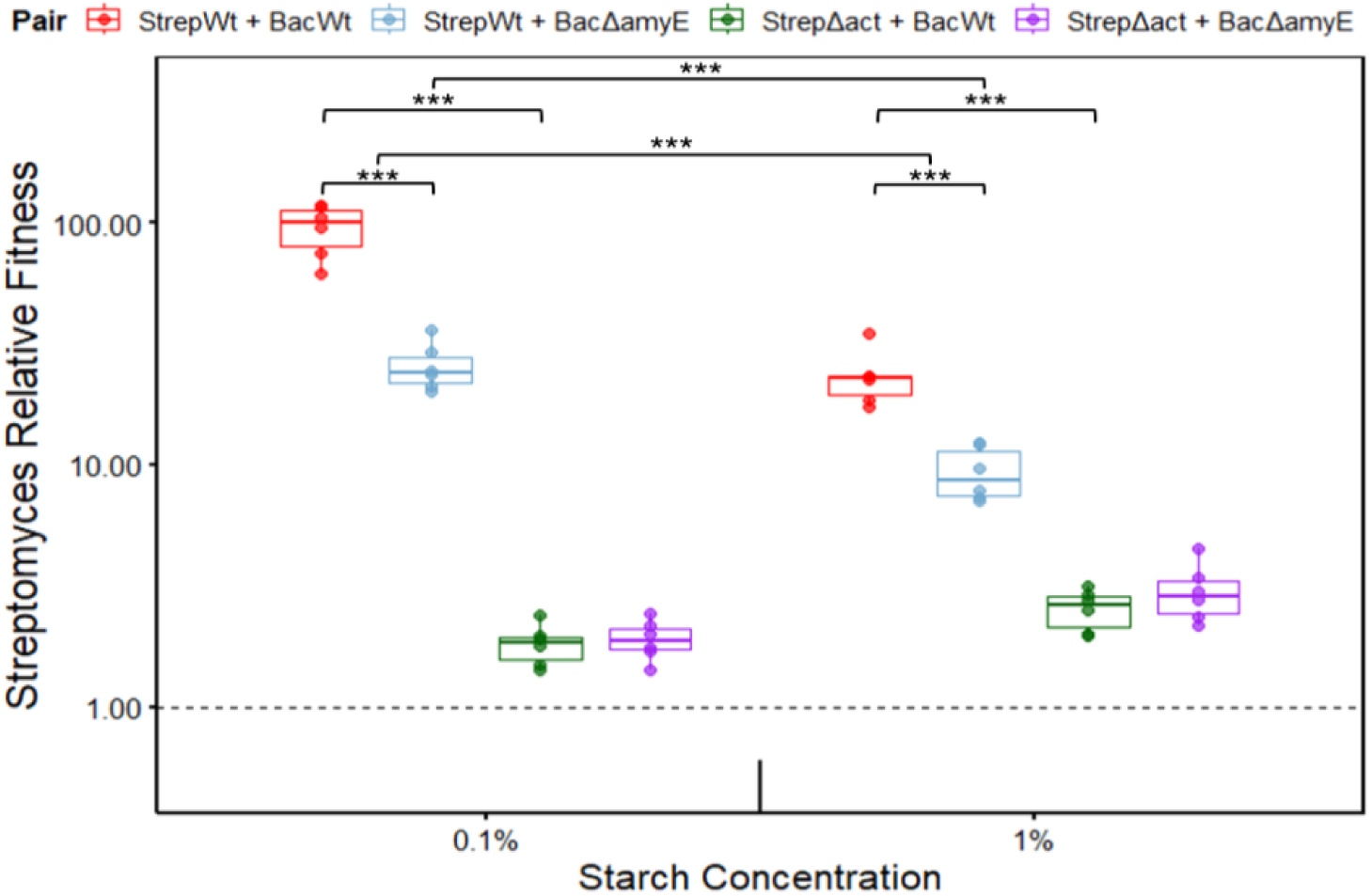
Fitness of *S. coelicolor* in competition with *B. subtilis* on 0.1% and 1% starch. On lower starch concentration, the benefit of producing antibiotics is greater, as is the fitness impact of resource exploitation. 1-way and 2-way ANOVAs were used to test statistical significance within and between starch concentration groups, respectively. *** corresponds to p < 0.001.

### Antibiotic production and colony growth trade-off at different starch concentrations

Differences in the fitness effects of antibiotic production or resource exploitation at different starch concentrations could be mediated by changes in antibiotic production, the growth of each strain, or some interaction between the two factors. To test this, we used halo assays to quantify antibiotic production of both *S. coelicolor* strains at 0.1% and 1% starch, while also measuring growth rate and spore production under the same conditions. As expected, the StrepΔact strain produces only negligible amounts of antibiotics (Figure 4A). By contrast, the WT strain produces significantly larger inhibition zones than the mutant regardless of starch concentration (ANOVA 2-way: F_1,20_ = 1750.58, p < 0.001). Additionally, StrepWt produces significantly larger inhibition zones when grown on lower starch concentrations (ANOVA 1-way: F_1,10_ = 63.47, p < 0.001). Results for growth rate and spore production (Figure 4B and 4C, respectively) are concordant and reveal the trade-off between antibiotic production and growth and sporulation. For both traits, we found that the StrepΔact strain grew more rapidly and produced more spores than the WT, regardless of the starch concentration (Growth Speed: ANOVA 2-way: F_1,561_ = 76.855, p < 0.001 / Colony CFU: ANOVA 2-way: F_1,44_ = 11.93, p < 0.001). In addition, growth rate and spore production were significantly higher for both strains when grown on 1% starch (Growth Speed: ANOVA 2-way: F_1,561_ = 37.001, p < 0.001; Colony CFU: ANOVA 2-way: F_1,44_ = 83.79, p < 0.001). Finally, we found a significant interaction between strain and starch concentration for growth speed (ANOVA 2-way Interaction: F_1,561_ = 9.295, p < 0.01), indicating that StrepWt was more affected by the increased resource concentration than StrepΔact. Taken together, these results suggest that antibiotic production comes at the expense of growth and that the shape of this relationship depends on resource availability.

**Figure 4.**
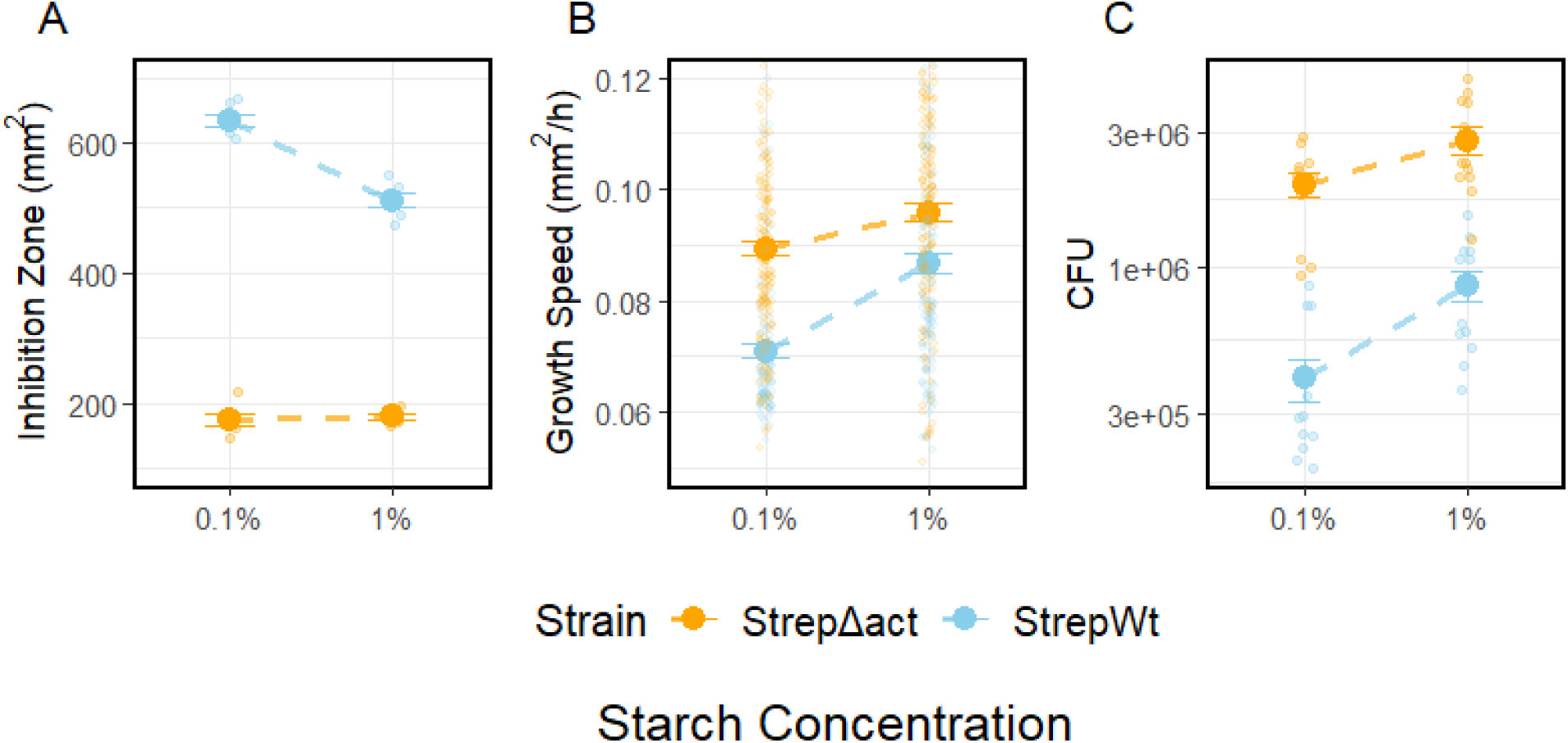
Inhibition, growth, and spore output of *S. coelicolor* on 0,1% and 1% starch. A) *Streptomyces* antibiotic inhibition halos of *B. subtilis* B) growth speed in (*mm*^2^/h) of individual colonies over 7 days of incubation. C) spore production after 7 days of incubation. Values correspond to means, and error bars correspond to SEM.

## Discussion

Microbes use different strategies to gain and retain access to resources and space in heterogeneous soil environments (Hibbing et al., 2010). While diffusible carbon can be directly consumed by most bacterial species, complex carbohydrates need to first be degraded by extracellular enzymes before they can be used (Wang & Kuzyakov, 2024). A consequence of extracellular metabolism is that resources produced by exoenzymes can be consumed by other bacteria, irrespective of whether or not they produce exoenzymes themselves (Allison, 2005). In monocultures, which are often used to study resource preferences or antibiotic production in the laboratory, this conflict never arises (Jauri et al., 2013; Romero et al., 2011)). However, in environments with higher biodiversity, the fitness of primary degraders can potentially be reduced, unless there are mechanisms to privatize resources by restricting competitor access (Sichert & Cordero, 2021; Smith & Schuster, 2019). Here, we sought to evaluate the fitness costs of exoenzyme exploitation during competitive interactions between species from two common soil genera, *Streptomyces* and *Bacillus*. We found that while resource exploitation reduced *S. coelicolor* fitness on complex resources, these costs could be significantly reduced if *S. coelicolor* was able to produce antibiotics. Moreover, we found that the benefits of antibiotics, and correspondingly the costs of resource exploitation, were higher in environments with lower resource concentrations. These results suggest that, despite their associated fitness costs, antibiotics can privatize public goods and support the idea that these compounds are used to protect resources for growth, rather than to protect resources released during development after growth has been arrested.

Given the ubiquity of resource exploitation, bacteria have evolved several different mechanisms to protect and privatize public goods (Smith & Schuster, 2019). Spatially structured populations, growth in biofilms, or physical attachment to insoluble complex carbohydrates can passively limit the diffusion of exoenzymes or degradation products (Drescher et al., 2014). High local cell densities and quorum sensing, coupled with spatial segregation, can restrict access of secreted public goods to clonemates, and this can be reinforced by mechanisms that promote cooperation between kin or that inhibit non-clonemates via interference competition (Cordero & Datta, 2016; Nadell et al., 2016). Although we are unaware of explicit mechanisms of kin recognition in streptomycetes, our results suggest that antibiotic production, especially when coupled to exoenzyme secretion (Nazari et al., 2013), could work as a type of active policing against resource exploitation by killing susceptible competitors to create competitor-free space. This use would be consistent with results showing that the benefits of antibiotic production are significantly higher in spatially structured environments (Westhoff S. et al. 2020) and supports the notion that antibiotics may be particularly important for slowly growing saprotrophs that specialize on patchily distributed insoluble resources. In well-mixed environments, by contrast, antibiotics may offer weaker benefits because both exoenzymes, antibiotics, and degradation products can diffuse away from producing cells or become too diluted (Chao & Levin, 1981). Indeed, recent results in *S. venezuelae* grown in liquid cultures showed that the products of chitin degradation were privatized by uptake through membrane transporters that effectively limited access to other competitors (Meij et al., 2025). Such “selfish” strategies have been observed in other primary degraders to restrict access to resource cheaters (Reintjes et al., 2020).

In addition to finding resource-dependent benefits of actinorhodin production in *S. coelicolor*, we also observed that the fitness benefits of antibiotic production were higher when competing against *B. subtilis* on lower starch concentrations. The fitness cost of resource cheating was similarly higher under these conditions. Cooperative traits are metabolically costly, and several studies have found that these costs decline as resource availability increases (Connelly et al., 2017). For example, *Pseudomonas aeruginosa* invested more heavily in cooperative biofilm exopolysaccharides and siderophores when grown with higher resource supply, likely because the costs of producing these products decline when resources are abundant (Brockhurst et al., 2008). Reciprocally, because the costs of cooperation are higher when resources are scarce, this implies that the fitness effects of cheating are correspondingly higher. While our results are consistent with these studies, they are also complicated by the fact that *S. coelicolor* fitness is due to the joint effects of antibiotic and exoenzyme production in our experiments. In low concentrations of starch, we hypothesize that the higher costs of resource cheating derive from increased reliance of both competitors on the products of starch degradation. Because *B. subtilis* grows more rapidly on these degradation products, this results in a stronger burden on *S. coelicolor* and therefore a higher cost of cheating. By contrast, the effects of cheating are diminished during growth on higher starch concentrations because the relative costs of amylase secretion are lower, while the benefits of amylase secretion show diminishing returns. Countering the negative effects of cheating, we observed significantly higher production of actinorhodin in lower concentrations of starch, leading to increased benefits. Although this also came at the expense of growth and sporulation, the benefits of antibiotic production that target *Bacillus* cheaters more than offset the effects of the trade-off.

Although we can only speculate about the mechanism underlying higher antibiotic production in lower concentrations of starch, it is known that *Streptomyces* antibiotics are in part regulated by resource type and availability (Ruiz-Villafán et al., 2022; Sánchez et al., 2010). For example, high concentrations of glucose and glycerol downregulate antibiotic formation (Romero-Rodríguez et al., 2018), while N-acetylglucosamine, a breakdown product of chitin, can induce actinorhodin production in *S. coelicolor* (Rigali et al., 2008). The response to N-acetylglucosamine is moreover strongly dependent on concentration, which is proposed to inform cells of either abundant resources for growth or impending famine. While this response can be understood intuitively as a mechanism by which *Streptomyces* optimizes the balance between growth, development, cell lysis, and defence, it is unclear if similar behaviors are found for other complex carbohydrates. Additionally, chitin-dependent production of antibiotics observed *in vitro* was more evident in soil microcosms, where growth on chitin induced both chitinases and secondary metabolites, including actinorhodin (Nazari et al., 2013). This suggests that the functional link between exoenzyme production and resource defense may in part be coordinated by unknown components of soil, in addition to the type or concentration of resource. Irrespective of the mechanism, our results confirm the sensitive interactions between resource availability and defensive strategies, and highlight the need for additional studies using a broader range of natural substrates, especially complex polysaccharides, and diverse competitors.

Our results broaden the scope of the many functional roles of *Streptomyces* antibiotics. However, our central conclusion, that costly resource exploitation can be mitigated by antibiotic production, is limited in a few ways. First, while growth on petri plates provides more spatial structure than mixed tubes, diffusion through agar is still extremely rapid, which might serve to homogenize the distribution of resources, exoenzymes, and antibiotics (Westhoff et al., 2020). Soil, by contrast, is a physically and compositionally heterogeneous environment where the transport and distribution of secreted (or generated) metabolites is influenced by soil hydration, particle size, adhesion, and the patchy distribution of insoluble complex carbohydrates (Wolf et al., 2013). While some of these factors could conceivably increase the benefits of antibiotics by creating higher local cell concentrations that drive inhibition, they could also lower antibiotic concentrations if production depends on high cell densities driven by quorum sensing. A second limitation concerns the fact that our experiments only examine interactions between two species. Although this helped to focus attention on extremes of antibiotic production and resource exploitation, it also neglects the potential impacts of higher-order strain interactions (both positive and antagonistic) or more complex cross-feeding dynamics in trophic cascades (van der Meij et al., 2017). Related to this, our results were limited to interactions at uniformly high cell densities and initially similar competitor frequencies, both of which are known to affect the local concentrations of antibiotics or exoenzymes, or the fitness of resource cheaters (van Gestel et al., 2014). Finally, it remains unclear how our results will generalize to other complex carbohydrates, especially if the costs of exoenzymes vary and if there are resource and concentration-dependent factors that coordinate antibiotic production with enzyme secretion or other aspects of growth and development. We will aim to address these limitations in subsequent studies.

## Acknowledgements

We thank NWO for funding and gratefully acknowledge Ákos T. Kovács for providing *Bacillus subtilis* strains and advice, Joost Willemse and Gerda Lamers help with microscopy and Matt Hutchings and Paul Hoskisson for comments on the manuscript.

## Supplementary Materials

### Bacillus growth curves on single carbon source

Bacterial growth was measured in Minimal Media (MM) which contains per litter: 0.5 g K2HPO4 (Avantor Performance Materials Poland S.A., Gilwice, Poland), 0.2 g MgSO4 (Merck KGaA, Darmstadt, Germany), 0.01 g FeSO4 (SIGMA-ALDRICH Co., St. Louis, USA). The solution pH was adjusted to 7.2 and amended with either 1% starch or 1% maltose. To measure the growth of *B. subtilis ΔamyE* on starch or maltose, we inoculated 0.1 OD of an overnight culture into wells of a 48-well plate containing 1 mL of media. A third negative control was included. OD600 was measured every 5 minutes under constant shaking (200 rpm) and 30 °C in a BioTek Epoch 2 Microplate reader (Agilent Technologies, USA).

### Mutant Comparison Halo of Inhibition

To compare the antibiotic activity of *S. coelicolor* antibiotic-producing mutants against *B. subtilis* (BacWt), we examined growth inhibition halos. Strains StrepΔact, StrepΔred, and StrepΔactΔred were all obtained from *S. coelicolor* A3(2) M145 modification (Floriano & Bibb, 1996). Undecylprodigiosin and actinorhodin production are encoded by the *red* and *act* gene, respectively. *Streptomyces* colonies were established by plating 2 µL drops containing 10^5^ *S. coelicolor* spores onto Minimal Media (MM) amended with 1% maltose. After five days of incubation, overnight cultures of *B. subtilis* Wt were diluted to roughly 10^8^/ml after which 100 µL was mixed with 4 mL molten LB soft agar (0.7%), and poured over the inoculated plates to create a uniform overlay lawn of susceptible bacteria. After 24 hours of incubation, the plates were imaged with a Nikon D200.

### Colony Growth Speed in Maltose

*Streptomyces* growth rate as a function of carbon source concentration was quantified by measuring radial growth via time-lapse imaging of replicate colonies. 10 µL containing 50 spores of StrepWt or StrepΔact were spread on modified ISP4 containing 1% maltose. Plates were incubated at 30 °C in a Reshape Imaging Device (Reshape Biotech, Copenhagen, Denmark) for 7 days. For each strain, two replicate plates were imaged every hour, allowing over 100 isolated colonies per plate to have their area calculated every hour using the Discovery platform of Reshape Biotech. Finally, Microsoft Excel (Office 365) was used to convert colony area into the average hourly area growth speed (*mm*^2^/*h*) of each colony.

## Supplementary data

**Supplemental Figure 1.**
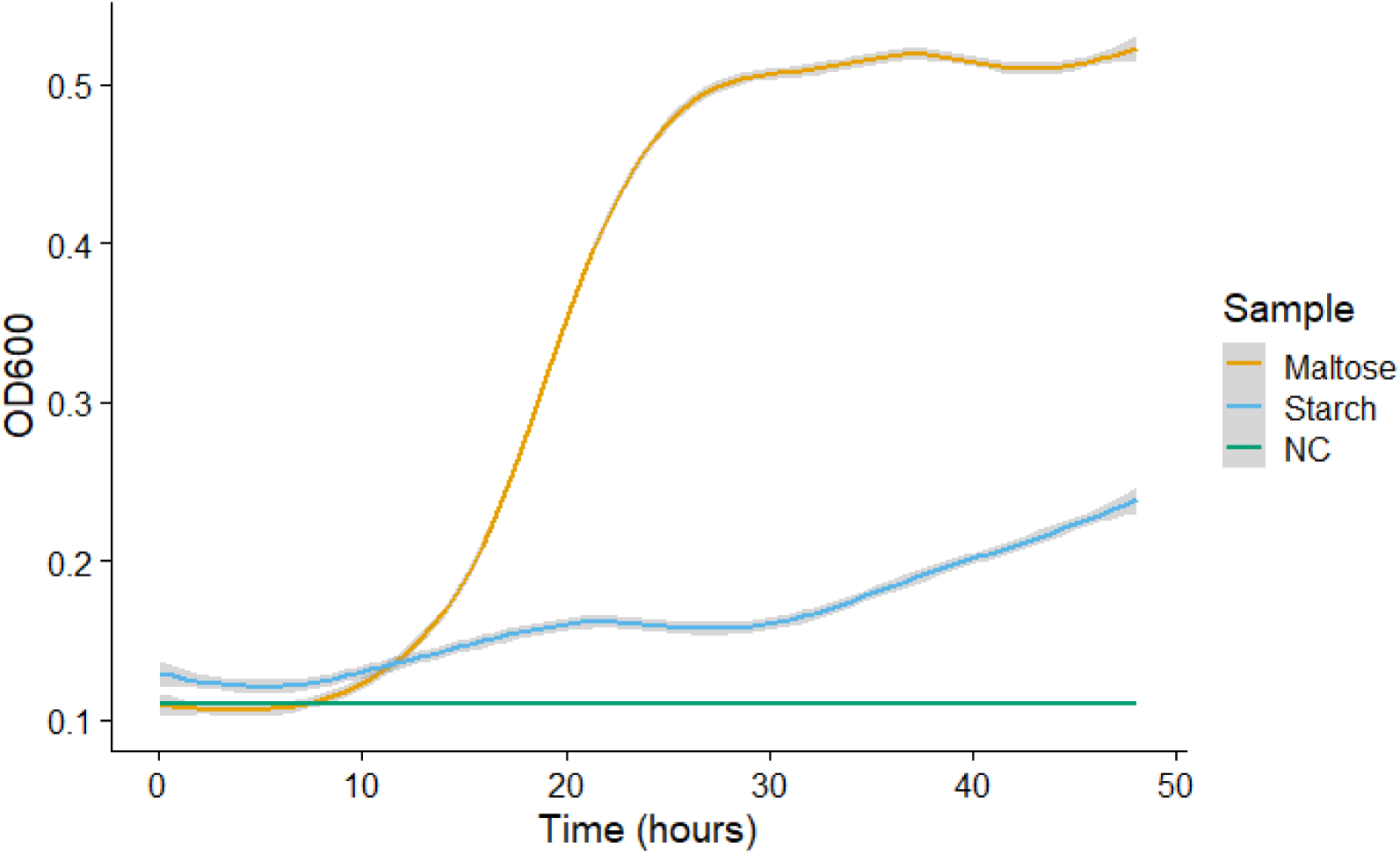
Growth curves of *B. subtilis ΔamyE* show amylase deficiency. *B. subtilis ΔamyE* was grown in liquid Minimal Media amended with either 1% starch or maltose in two technical replications. Lines were generated with the ggplot2 geom_smooth function.

**Supplemental Figure 2.**
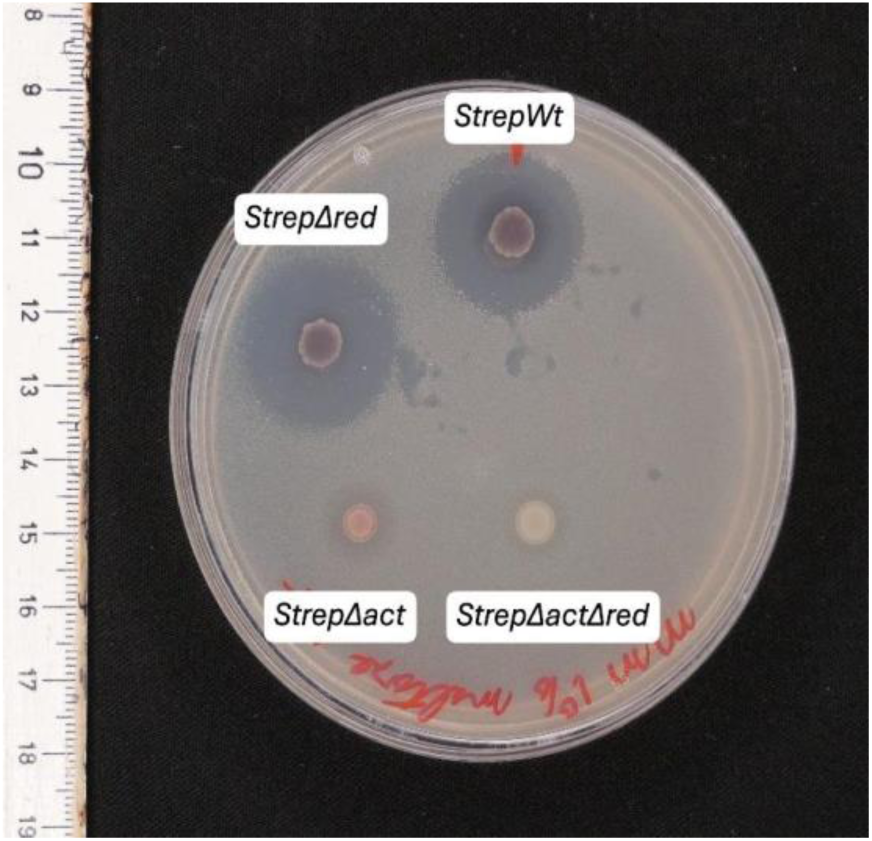
*S. coelicolor* inhibition against *B. subtilis*. Halo size depends on actinorhodin production, not undecylprodigiosin, encoded by the red and act gene, respectively. Antibiotic production was measured on Minimal Media amended with 1% maltose. A soft agar (0.7%) LB overlay with *B. subtilis* was poured after 2 days of sole *Streptomyces* incubation. Imaging occurred 24 hours after pouring the overlay.

**Supplemental Figure 3.**
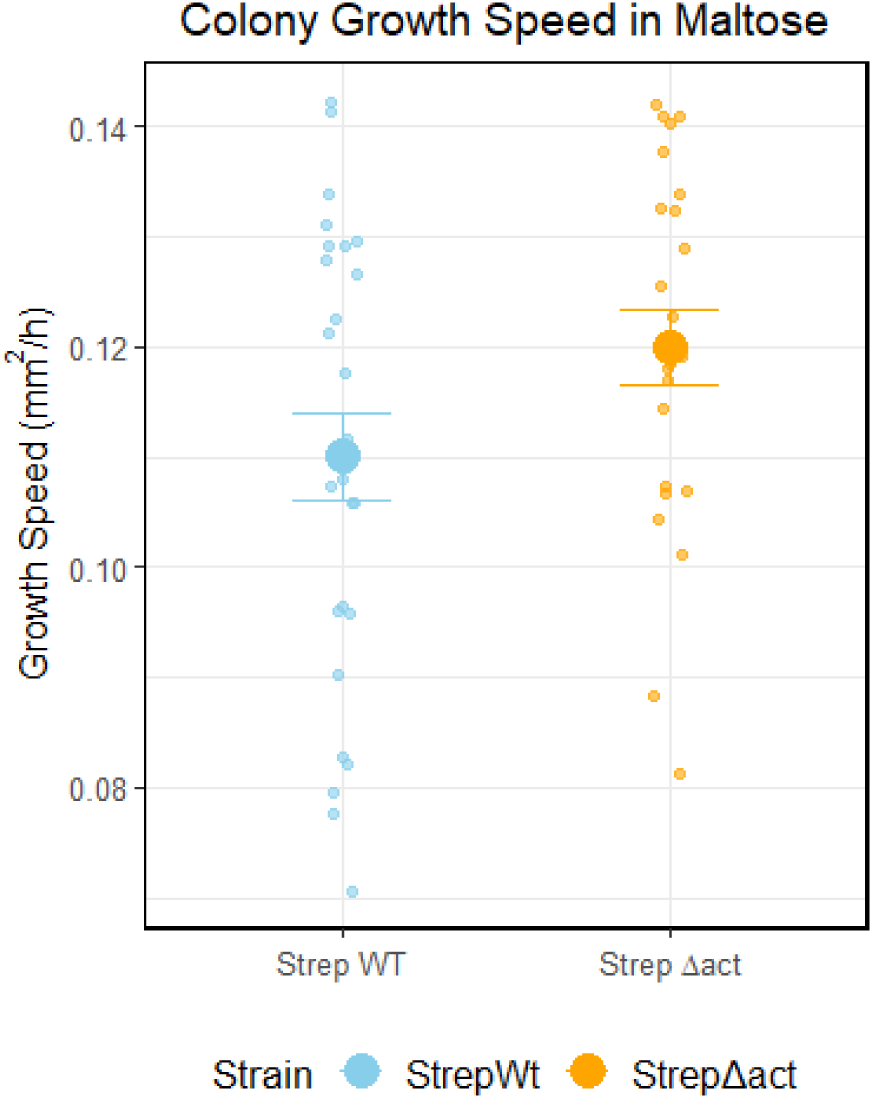
*S. coelicolor* growth speed on ISP4 amended with 1% maltose. Petri dishes with about 30 single colonies were tracked, and the average growth speed (*mm*^2^/h) of each colony was measured over 7 days using a Reshape Imaging Device. Growth speed was calculated by averaging the difference in area between every hour. Values correspond to means, while error bars correspond to the standard error of the mean.

